# Actin associates with actively elongating genes and binds directly to the Cdk9 subunit of P-TEFb

**DOI:** 10.1101/2023.09.06.556538

**Authors:** Salla Kyheröinen, Bina Prajapati, Maria Sokolova, Maximilian Schmitz, Tiina Viita, Matthias Geyer, Maria K. Vartiainen

## Abstract

Nuclear actin has been demonstrated to be essential for optimal transcription, but the molecular mechanisms and direct binding partner for actin in the RNA polymerase complex have remained unknown. By using purified proteins in a variety of biochemical assays, we demonstrate a direct and specific interaction between monomeric actin and Cdk9, the kinase subunit of the positive transcription elongation factor b (P-TEFb) required for RNA polymerase II pause-release. This interaction efficiently prevents actin polymerization, is not dependent on kinase activity of Cdk9 and is not involved with releasing P-TEFb from its inhibitor 7SK snRNP complex. Supporting the specific role for actin in the elongation phase of transcription, chromatin immunoprecipitation followed by deep sequencing (ChIP-seq) reveals that actin interacts with genes only upon their active transcription elongation. This study therefore provides novel insights into the mechanisms by which actin facilitates the transcription process.

## Introduction

Actin is an abundant protein having essential roles both in the cytoplasm and in the cell nucleus. It is a key component of the cytoskeleton, but inside the nucleus its functions are mostly related to gene expression control and maintenance of genomic integrity (Ulferts et al., 2021). The basic unit of actin is the globular monomer (G-actin) that can polymerize into filaments (F-actin) and the proper balance of these forms, maintained by actin-binding proteins (ABPs), is critical for actin functions. Shuttling of actin between the cytoplasmic and nuclear compartments is an active process and the two pools of actin are dynamically connected. Actin is imported into the nucleus by Importin-9 (Ipo9) together with small ABP cofilin (Dopie et al., 2012) and exported by Exportin-6 (Exp6) as a complex with ABP profilin (Stuven et al., 2003). Nuclear actin is an important regulator of gene expression. G-actin is a stoichiometric subunit of several chromatin remodeling and modifying complexes, which control the structure and accessibility of chromatin (Klages-Mundt et al., 2018). Actin also directly regulates the activity of specific transcription factors, such as myocardin-related transcription factors (MRTFs) that are co-activators of the serum response factor (SRF). G-actin binding to MRTF prevents SRF activation (Vartiainen et al., 2007), and there are various mechanisms to regulate MRTF/SRF pathway via altering actin nucleo-cytoplasmic transport and polymerization status (Sidorenko and Vartiainen, 2019).

One of the essential functions of nuclear actin is its role in transcription regulation. For optimal transcription, the levels of actin inside the nucleus have to be controlled, as manipulating actin levels impairs transcription both *in vitro* and in cells (Dopie et al., 2012; Hu et al., 2004; Philimonenko et al., 2004; Sokolova et al., 2018). Decreased nuclear actin levels in response to laminin-111 and consequent PI3 kinase pathway attenuation leading to increased Exp6 activity have also been shown to destabilize Pol II and Pol III interactions with nuclear substructures, causing lower RNA synthesis rates and quiescence in mouse epithelial cells (Fiore et al., 2017; Spencer et al., 2011). In *Drosophila* ovaries, actin and Pol II co-occupy promoter regions of most transcribed genes, as well as the gene bodies of highly transcribed genes (Sokolova et al., 2018). It is becoming clear that actin has functions throughout the transcription process. Actin has been linked to pre-initiation complex formation (Hofmann et al., 2004) and F-actin seems to cluster Pol II on inducible genes (Wei et al., 2020), promoting the formation of the transcription machinery. Actin also functions in the elongation phase and in Pol I mediated transcription, also nuclear myosin I is suggested to be involved, possibly combining the motor function of myosin with actin polymerization (Almuzzaini et al., 2016; Philimonenko et al., 2004; Ye et al., 2008). Furthermore, actin has been linked to Pol II pause-release (Qi et al., 2011), an important control point of transcription by Pol II, in which the polymerase pauses shortly after elongation has started and needs positive transcription elongation factor b (P-TEFb) to proceed into productive elongation (Core and Adelman, 2019). In the transcription elongation phase, actin also associates with heterogeneous nuclear ribonucleoprotein U (hnRNP U) (Kukalev et al., 2005), and the function of actin here might be to recruit transcription coactivators, such as PCAF histone acetyltransferase (Obrdlik et al., 2008). Finally, actin takes part in pre-mRNA processing, as previous results from our lab show that manipulating actin disturbs alternative splicing, either through a direct role in splicing, or by affecting Pol II elongation or pausing rate so that also weaker splicing sites could be used (Viita et al., 2019). Altogether, there seems to be distinct roles for both monomeric and filamentous actin in transcription, and some interactions, in which the form of actin is not yet known.

Despite vast evidence of actin functions in transcription, the direct binding partner of actin within the RNA polymerases is not known. Actin has been suggested to co-purify with all three RNA polymerases (Pol): Pol I (Fomproix and Percipalle, 2004), Pol II (Egly et al., 1984; Smith et al., 1979) and Pol III (Hu et al., 2004) and was already in the 1980’s indicated to be involved in transcription of protein coding genes (Scheer et al., 1984). However, the form of actin associating with the polymerases is not known and the interaction remains to be characterized at a molecular level. Our mass spectrometry screen for nuclear actin interactome contained several transcription-related proteins as hits with BioID technique, detecting more transient interactions, but not in AP-MS that is best suited for stable binding events (Viita et al., 2019). Also, most of the same proteins were captured with an actin mutant incapable of polymerization, suggesting that F-actin formation would not be needed for these interactions to take place. One of the BioID hits was both subunits of the P-TEFb complex (Viita et al., 2019). Here we demonstrate that actin, in its monomeric form, binds directly and specifically to the P-TEFb subunit Cdk9. We further demonstrate that actin associates with genes only upon their active elongation, and that the kinase activity of Cdk9 is neither required for the interaction with actin, nor actin recruitment to the transcribing genes. Functionally, actin is not involved in releasing P-TEFb from its inhibitory complex, and modestly influences Cdk9 kinase activity.

## Results

### Actin is recruited to the genes only upon active transcription elongation and does not directly interact with the RNA polymerase II

Previous studies have implicated actin in several steps of transcription (Hofmann et al., 2004; Kukalev et al., 2005; Obrdlik et al., 2008; Qi et al., 2011; Viita et al., 2019). By using chromatin immunoprecipitation followed by deep-sequencing (ChIP-seq), we have demonstrated that actin interacts with all transcribed genes near transcription start sites (TSS) in *Drosophila melanogaster* ovaries. In addition, we showed that actin accumulates on gene bodies of highly transcribed genes required for egg shell formation during *Drosophila* oogenesis (Sokolova et al., 2018). Heat shock protein (*hsp)* genes encode chaperone proteins that are required to maintain proteostasis not only during proteotoxic stress, such as heat shock (HS), but also during normal conditions. Indeed, many *hsp* genes are expressed also during *Drosophila* oogenesis (Ambrosio and Schedl, 1984). To study, if actin is binding to *hsp* genes *in vivo*, we used ChIP-seq on fly ovaries at room temperature (25°C). Similarly as we have reported earlier for the *chorion* genes (Sokolova et al., 2018), actin was found on the gene body of *Hsp68* (Fig 1A). RNA polymerase II phosphorylated on serine 5 (Pol II), peaked near the transcription start site (TSS), but was also abundant on the gene body (Fig 1A). Analysis of the 15 *hsp* genes revealed similarly that Pol II is more concentrated near the TSS than actin, which displayed a rather broad distribution centered on the coding region of the gene (Fig 1B). Thus, actin and Pol II do not display identical binding patterns on the *hsp* genes.

**Figure 1.**
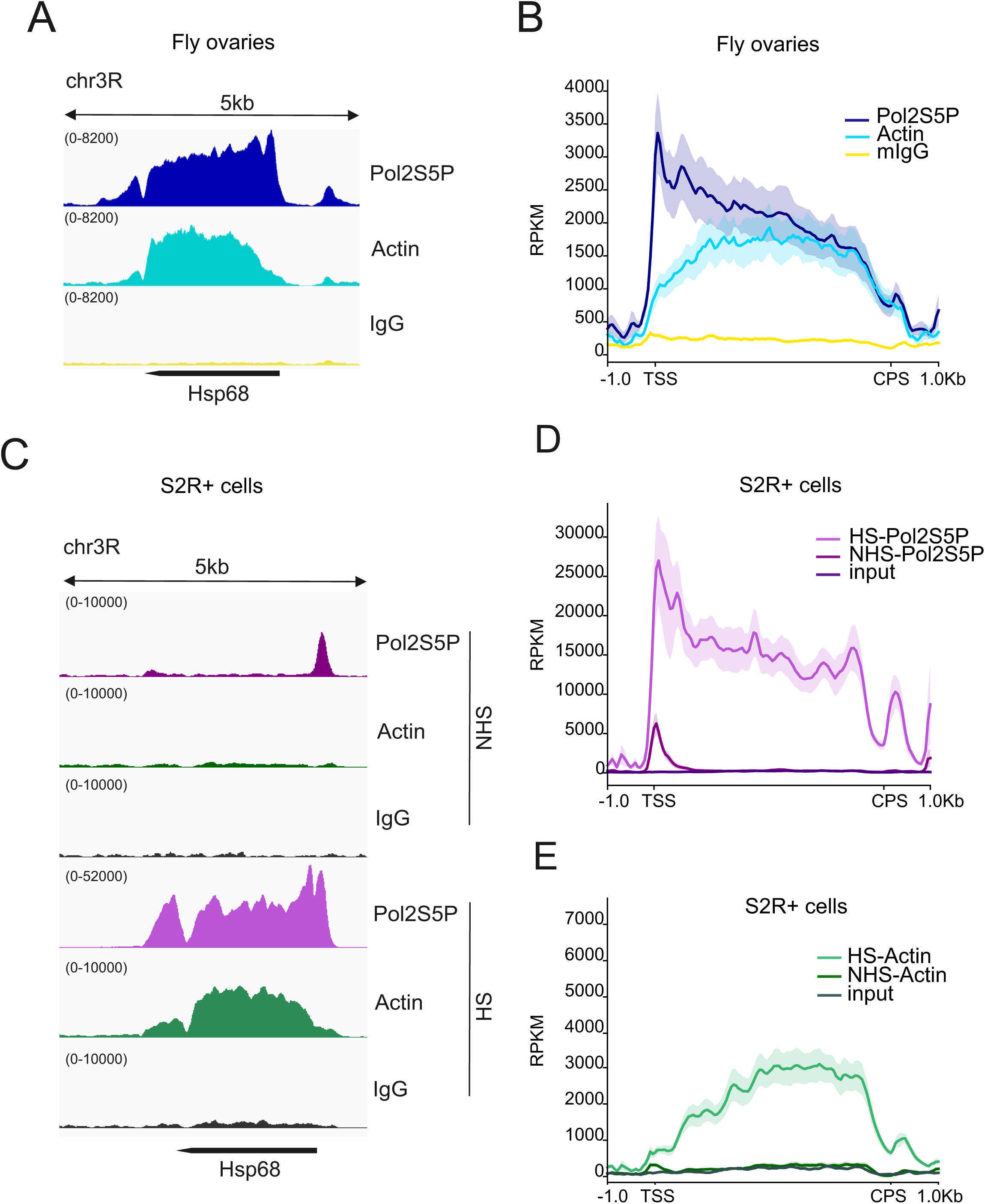
Actin associates with *hsp* genes only upon their active transcription elongation. A. Actin binds *hsp* genes in *Drosophila* ovaries. Normalized (RPKM) coverage of RNA polymerase phosphorylated on serine 5 (Pol2S5P), actin and normal IgG from chromatin immunoprecipitation followed by deep sequencing (ChIP-seq) on *Hsp68* in *Drosophila* ovaries (W1118). B. Pol II S5P, actin and mouse IgG coverage on 15 heat shock responding genes in *Drosophila* ovaries. Normalized fragment counts are scaled to 5 kb gene size. Transcription start site (TSS) and cleavage and polyadenylation site (CPS). C. Actin binds to *hsp* genes only upon their transcriptional activation during HS. Normalized (RPKM) coverage of Pol II S5P, actin and normal IgG on Hsp68 from ChIP-seq in S2R+ cells in non heat shock (NHS) and heat shock (HS, 20 min) conditions. D. Pol II S5P coverage on 15 heat shock responding genes in S2R+ cells in non heat shock (NHS) and heat shock (HS) conditions. Input is used as a negative control and data shown as in B. E. Actin coverage on 15 heat shock responding genes in S2R+ cells in non heat shock (NHS) and heat shock (HS) conditions, shown as in D.

Since *hsp* genes are already abundantly expressed in fly ovaries without HS, we turned to the cultured fly cell line S2R+ as a model to study robust HS-induced transcriptional activation, and performed ChIP-seq of actin and Pol II in cells cultured at room temperature (non-heat shock; NHS) or heat shocked for 20 min at 37°C (Fig 1C-E). In NHS conditions, Pol II is found mainly paused near the TSS of *Hsp68* gene (Fig 1C). However, actin did not display any significant enrichment on the *Hsp68* gene (Fig 1C) under these conditions, when the gene is not being actively transcribed. When the cells were heat shocked, Pol II was strongly recruited to the *Hsp68* gene and similarly as in fly ovaries, the distribution peaked near the TSS. Nevertheless, enrichment at the gene body and after the cleavage and polyadenylation site (CPS) was also very evident (Fig 1C). Upon HS, and similarly as observed in fly ovaries, actin was enriched on the coding region of *Hsp68*, but did not show particular enrichment near TSS (Fig 1C). Analysis of the 15 *hsp* genes revealed the same pattern (Fig 1D-E). Actin is not significantly associated with the *hsp* genes prior to their transcriptional activation by HS, but only accumulates to the genes upon their active transcription elongation (Fig 1E). The binding patterns of Pol II and actin are not similar (Fig 1D-E), indicating that actin does not interact with all Pol II complexes.

In several earlier studies, actin has been shown to co-purify with the RNA polymerases (Egly et al., 1984; Fomproix and Percipalle, 2004; Hu et al., 2004; Smith et al., 1979), but the details of this interaction have remained uncharacterized. We purified whole, transcriptionally active human Pol II from HeLa nuclear pellets to gain further insight into the possible interaction. However, in our Western blot of samples taken during the purification process (Fig S1A), actin was present in the first steps of the purification, but was lost during anion exchange (samples 3-4), indicating that the interaction is not stable and that actin is not an integral component of the purified polymerase, since it is not found in the final protein preparation (Fig S1A, sample 8). We then studied, if the purified proteins would form a complex that would persist in size-exclusion chromatography (SEC) using buffer conditions optimal for actin and NaCl concentration of 100 mM to improve the performance of the gel filtration column. In this experiment, actin was complexed with Latrunculin B (LatB), a small molecule inhibiting actin polymerization that would otherwise take place when salt is present. The two proteins eluted from the SEC column in different fractions (Fig S1B), which means that no complex was formed. This experiment suggests that actin, in its monomeric form, cannot directly interact with Pol II. This finding is further supported by our mass spectrometry (MS) screen for nuclear actin binding partners, which did not identify any Pol II subunits as high-confidence interactors in either affinity-purification-MS or BioID experiments (Viita et al., 2019).

### Actin monomers bind directly to the Cdk9 subunit of P-TEFb

Our ChIP-seq data demonstrated that actin bound only to actively elongating genes (Fig 1C, E), suggesting that a protein specifically required for transcription elongation may recruit actin to the transcribing polymerase. Previous studies had demonstrated that actin can be co-immunoprecipitated with P-TEFb (Qi et al., 2011), which plays a critical role in releasing the paused polymerase to active transcription elongation. Moreover, we had identified P-TEFb as a high-confidence interactor in our MS-based nuclear actin interactome analysis (Viita et al., 2019). Nevertheless, neither of these studies were able to establish whether the interaction between actin and P-TEFb is direct or not. To study this further, we first confirmed the interaction between these two proteins by using co-immunoprecipitation assay. Indeed, Flag-tagged Cdk9, the kinase subunit of P-TEFb, could be co-immunoprecipitated with HA-tagged actin, but not with an empty vector control (Fig 2A). To gain insight into the functional form of actin involved in this interaction, we then performed the experiment using actin mutants that are either predominantly monomeric (R62D, G13R) (Posern et al., 2002), favor polymerization (V159N) (Posern et al., 2002) or have mutations in the hydrophobic cleft, which is a commonly used site for actin-binding in many proteins (Cao et al., 2016). Rather surprisingly, all tested actin mutants were able to co-immunoprecipitate Cdk9, and the amount of Cdk9 detected in the IP sample seemed to correspond to the expression levels of Cdk9 construct in the input sample (Fig 2B). Thus, this experiment did not provide further insights into the form of actin engaged with P-TEFb.

**Figure 2.**
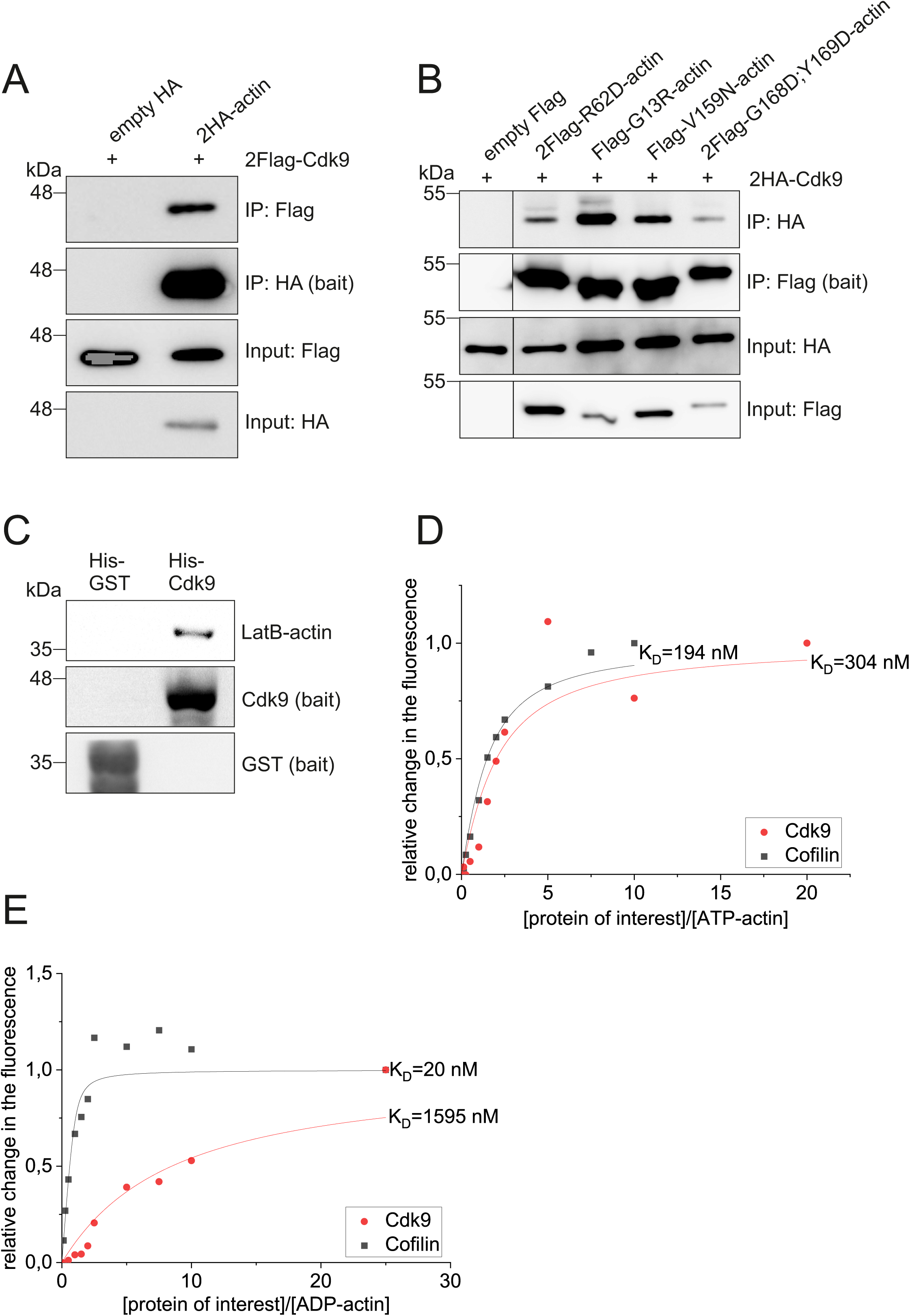
Cdk9 binds directly to G-actin with a preference towards ATP-actin. A. Western blot of co-immunoprecipitation (IP) experiment for 2Flag-Cdk9 (prey) and 2HA-actin (bait), using empty HA as a control. B. Western blot of co-IP experiment for 2HA-Cdk9 (prey) and indicated Flag/2Flag-tagged actin mutants (baits). Boxed areas cut from the same exposure to remove a construct with weak expression level and a lane with protein ladder. C. Western blot of pulldown assay of LatB-actin using His-Cdk9 as the bait and His-GST as a control bait. His-GST detected with His-HRP antibody, His-Cdk9 with Cdk9 antibody and LatB-actin with Ac-40 actin antibody. D. NBD-actin monomer-binding assay showing increase in the fluorescence of 0.2 μM NBD-labeled ATP-Mg-G-actin under physiological ionic conditions over a range of concentrations of Cdk9 or known actin binding protein cofilin. Symbols are data points and lines represent the exponentially fitted binding curves. N=3, one example presented. Experimentally determined dissociation constants (K_D_) calculated according to formula presented in materials and methods. Average K_D_ of three assays is 375 nM. E. NBD-actin monomer binding assay as in A, but using ADP-bound actin.

To study, if the binding between actin and P-TEFb is direct, we purified His-tagged Cdk9 from insect cells and actin from rabbit skeletal muscle. We then complexed the actin with Latrunculin B (LatB-actin) to perform a pulldown assay using His-Cdk9 as a bait under physiological salt conditions. LatB-actin was detected on beads coated with His-Cdk9, but not on beads with His-GST lysate (Fig 2C). This assay demonstrates a direct interaction between actin and the Cdk9 subunit of P-TEFb, and also indicates that actin interacts with P-TEFb as a monomer.

Next, we used the NBD actin monomer-binding assay (Fig 2D-E) to characterize the interaction between actin and Cdk9 further. In this assay, lysine 372 of actin is labeled with NBD fluorophore and upon protein binding near this residue, fluorescence of NBD changes and the changes can be recorded (Detmers et al., 1981). Actin is always bound to a nucleotide, ATP or ADP, and this is a critical factor that controls the polymerization process and interactions with other proteins. In addition, actin secondary structure is different between ATP-and ADP-bound forms (Zheng et al., 2007). Many actin monomer-binding proteins, such as thymosin β4, profilin and MIM prefer to bind ATP-actin (Carlier et al., 1993; Mattila et al., 2003; Vinson et al., 1998), whereas cofilin and twinfilin have higher affinity towards ADP-actin (Carlier et al., 1997; Ojala et al., 2002). With the NBD-actin assay, we observed that Cdk9 has a higher affinity towards ATP-actin (Fig 2D) compared to ADP-actin (Fig 2E). Average experimentally defined dissociation constant (K_D_) towards ATP-actin from three independent replicate experiments was 375 nM, but for ADP-actin the K_D_ could not be reliably defined, since the affinity was low. Cofilin was used as a control in our experiment and the K_D_ values obtained experimentally here for cofilin binding match earlier results (K_D_ = 20-100 nM towards ADP-actin) (Blanchoin and Pollard, 1998; Carlier et al., 1997; Vartiainen et al., 2002), verifying that the assay worked as expected. The NBD-actin assay thus further confirmed the direct interaction between actin monomers and Cdk9, and suggested that the interaction is sensitive to the nucleotide status of actin.

### P-TEFb inhibits actin polymerization and the interaction is not dependent on the kinase activity of the complex

To study how binding of P-TEFb affects actin, we next investigated actin polymerization with purified proteins by using pyrene actin polymerization assay (Fig 3). In this assay, G-actin is labeled with fluorophore pyrene and buffer conditions are changed to favor actin polymerization, which can be measured by following the increase in pyrene fluorescence as filaments form (Doolittle et al., 2013). The P-TEFb complex used in these assays consisted of full-length Cdk9 and a C-terminally truncated version of Cyclin T1 (amino acids 1-272) containing the cyclin-boxes only. This N-terminal domain of Cyclin T1 is widely used in both P-TEFb crystal structures (Baumli et al., 2008) and *in vitro* biochemical experiments, and demonstrated to activate Cdk9 (Bieniasz et al., 1999).

**Figure 3.**
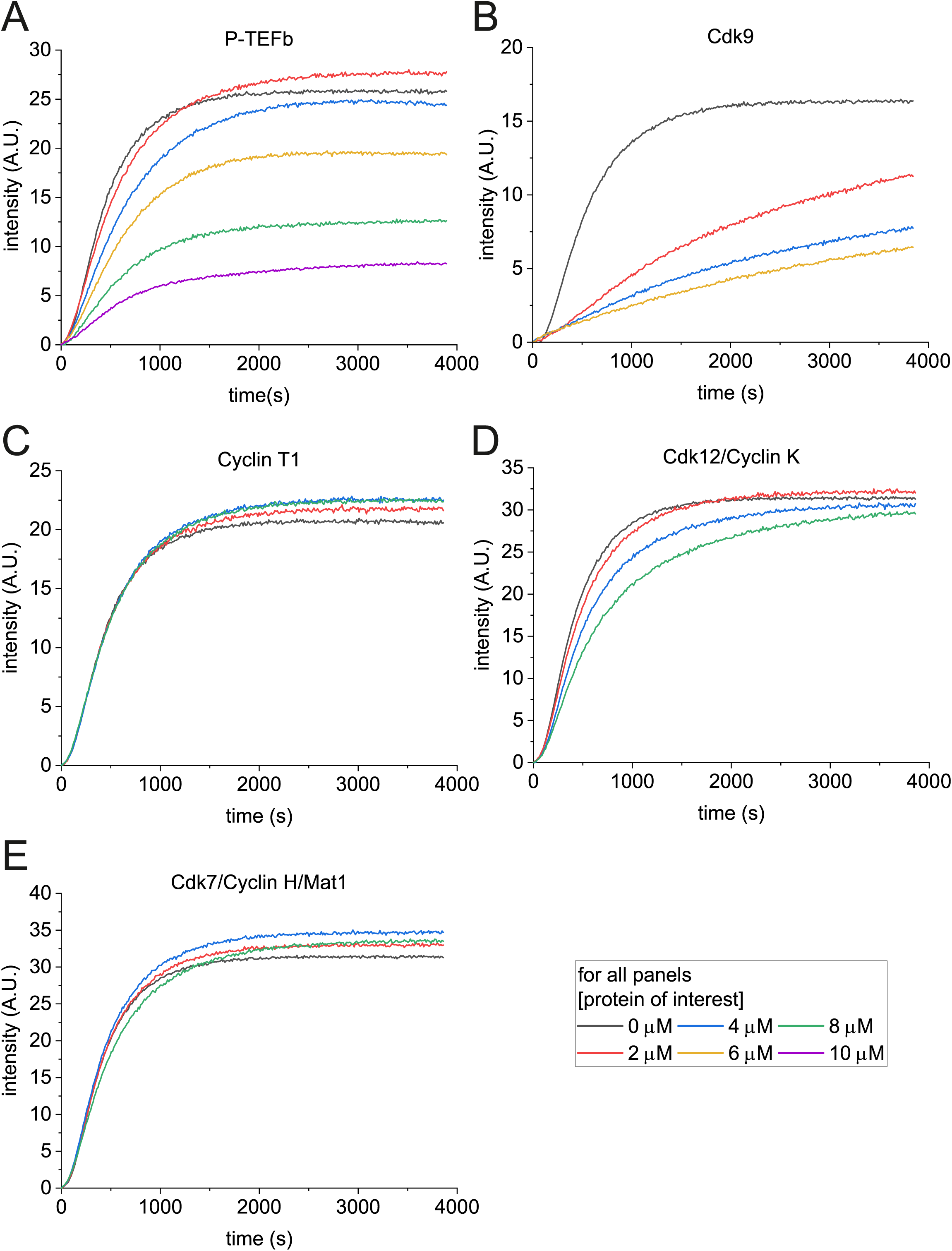
Actin binds to P-TEFb via Cdk9 and the binding inhibits actin polymerization. Pyrene actin polymerization assay with A. P-TEFb, B. Cdk9, C. Cyclin T1, D. Cdk12/Cyclin K, E. Cdk7/Cyclin H/Mat1 added in increasing concentrations and actin polymerization monitored by increase in the pyrene fluorescence. Actin concentration 4 μM, except for in the experiment with Cdk9 where actin is 2.5 μM. An example graph from one assay is presented.

Our results show that actin binding to P-TEFb (Fig 3A) and to Cdk9 alone (Fig 3B) inhibits actin polymerization in a concentration dependent manner, likely by sequestering actin monomers. However, Cyclin T1 alone did not have any effect on actin polymerization (Fig 3C), further confirming that Cdk9 is the binding partner for actin in the P-TEFb complex, as suggested already by our pulldown (Fig 2C) and NBD-actin assays with Cdk9 (Fig 2D). To test, if the interaction is specific to P-TEFb, or a more general phenomenon for transcriptional Cdk/cyclin complexes, we tested purified Cdk12/Cyclin K and Cdk7/Cyclin H/Mat1 complexes in the pyrene actin assay (Fig 3D-E). Cdk7/Cyclin H/Mat1 did not influence actin polymerization (Fig 3E), whereas Cdk12/Cyclin K modestly decreased actin polymerization (Fig 3D), but the effect was clearly weaker than observed with P-TEFb.

Since actin binds to the P-TEFb complex via Cdk9, which is a kinase, we wondered whether the kinase activity of Cdk9 might be required for the interaction. To this end, we used Cdk9 inhibitors flavopiridol (Chao et al., 2000), NVP-2 (Olson et al., 2018) and iCdk9 (Lu et al., 2015) in the pyrene actin polymerization assay. None of the inhibitors alone influenced actin polymerization (Fig 4A). Moreover, they also did not affect the ability of Cdk9 to inhibit actin polymerization (Fig 4A), leading us to conclude that the kinase activity is not needed for Cdk9 binding to actin monomers. To explore this further, we took advantage of the strong binding of actin to the gene bodies of *hsp* genes upon their transcription activation with HS (Fig 1C-E) and performed ChIP-qPCR experiments on the HS-responding gene *Hsp70*. Interestingly, and in line with the biochemical experiments (Fig 4B), inhibiting Cdk9 activity did not reduce actin-binding to the *Hsp70* gene. Thus, kinase activity of Cdk9 is dispensable for both actin binding to P-TEFb and to actin recruitment to actively transcribing genes.

**Figure 4.**
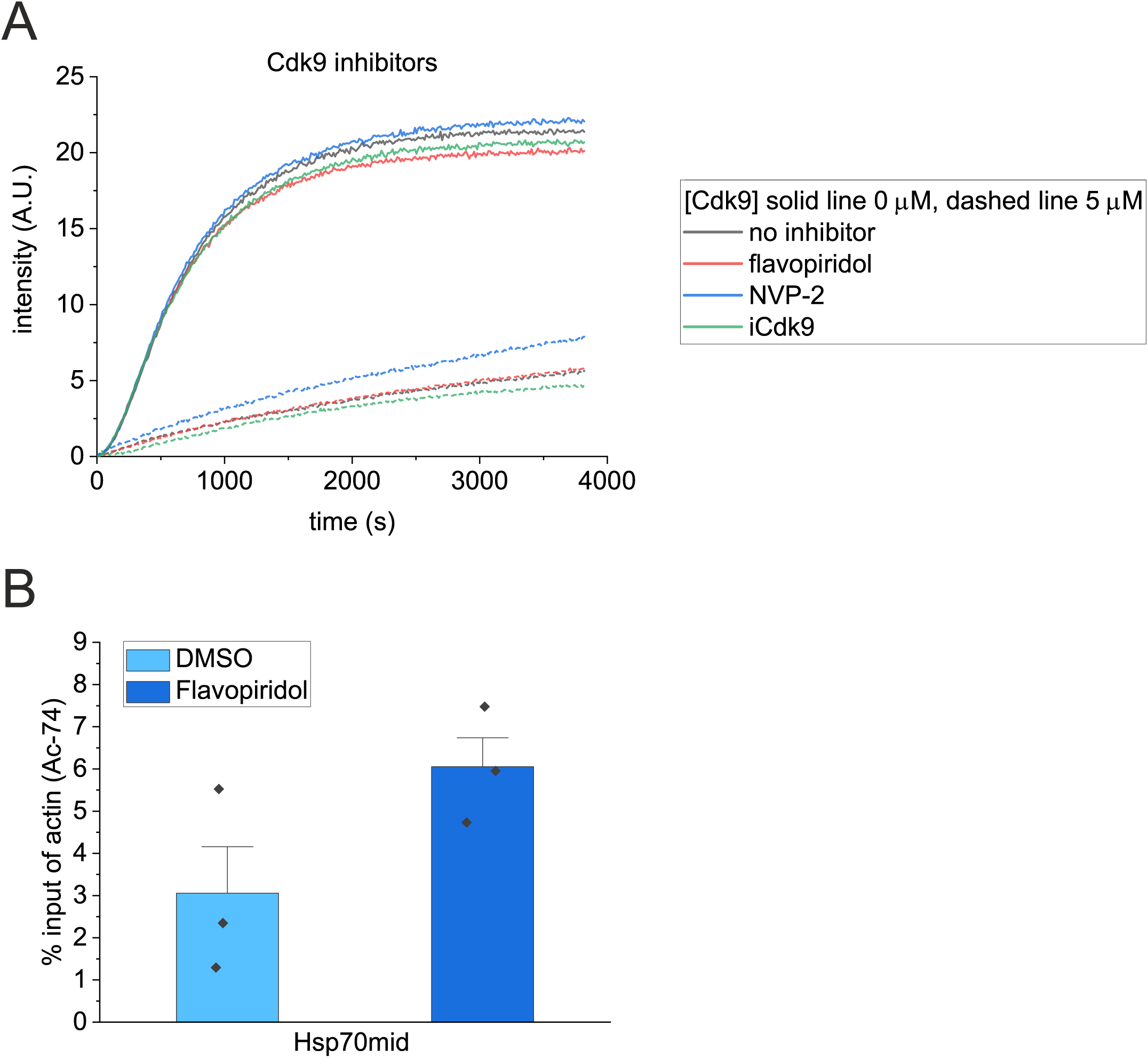
Kinase activity of Cdk9 is not required for the interaction with actin or actin recruitment to *hsp* genes. A. Pyrene actin polymerization assay with Cdk9 (0 and 5 μM) and Cdk9 inhibitors. Flavopiridol was 15 μM, NVP-2 5 μM and iCdk9 10 μM. Actin concentration 2.5 μM. B. Chromatin immunoprecipitation with actin Ac-74 antibody followed by RT-qPCR (ChIP-qPCR) of *hsp70* middle region (Hsp70mid) in *Drosophila* S2R+ cells. Cells were treated with DMSO/Flavopiridol for 10 min, followed by 20 min heat shock and immediate fixing. N=3, error bars ± 0.5 SD. Data presented as % of input enrichment. Statistical significance tested with two-sample t-test, DMSO vs. Flavopiridol P=0.130.

### Actin does not release P-TEFb from the inhibitory complex and has a minor influence on its kinase activity

In cells, most of P-TEFb is bound to an inhibitory 7SK snRNP complex that is needed to control P-TEFb kinase activity and consequently Pol II transcription (C. Quaresma et al., 2016). Apart from P-TEFb itself, our BioID assay for nuclear actin interacting proteins (Viita et al., 2019) identified P-TEFb regulators LARP7, BRD4 and DDX21 as hits. This led us to hypothesize that actin could control the association of P-TEFb with these factors or the release of P-TEFb from 7SK snRNP, since LARP7 is part of the inhibitory complex, binding P-TEFb to promote its incorporation into the 7SK snRNP and regulating the stability of the 7SK (C. Quaresma et al., 2016). To test this idea, we performed an *in vitro* P-TEFb release assay, in which 7SK snRNP from HeLa cells was immobilized with HEXIM1 antibody and purified muscle actin was added to the reaction as a possible release agent under conditions that keep actin monomeric. As expected, RNAse A released P-TEFb from the complex by degrading the 7SK RNA, and thus reduced the amount of Cdk9 that was co-immunoprecipitated with HEXIM1 (Fig 5A). However, actin, at two different concentrations, did not reduce Cdk9 in the HEXIM1 co-immunoprecipitation (Fig 5A). Thus, at least under these *in vitro* conditions, actin is not able to release P-TEFb from the inhibitory 7SK snRNP complex.

**Figure 5.**
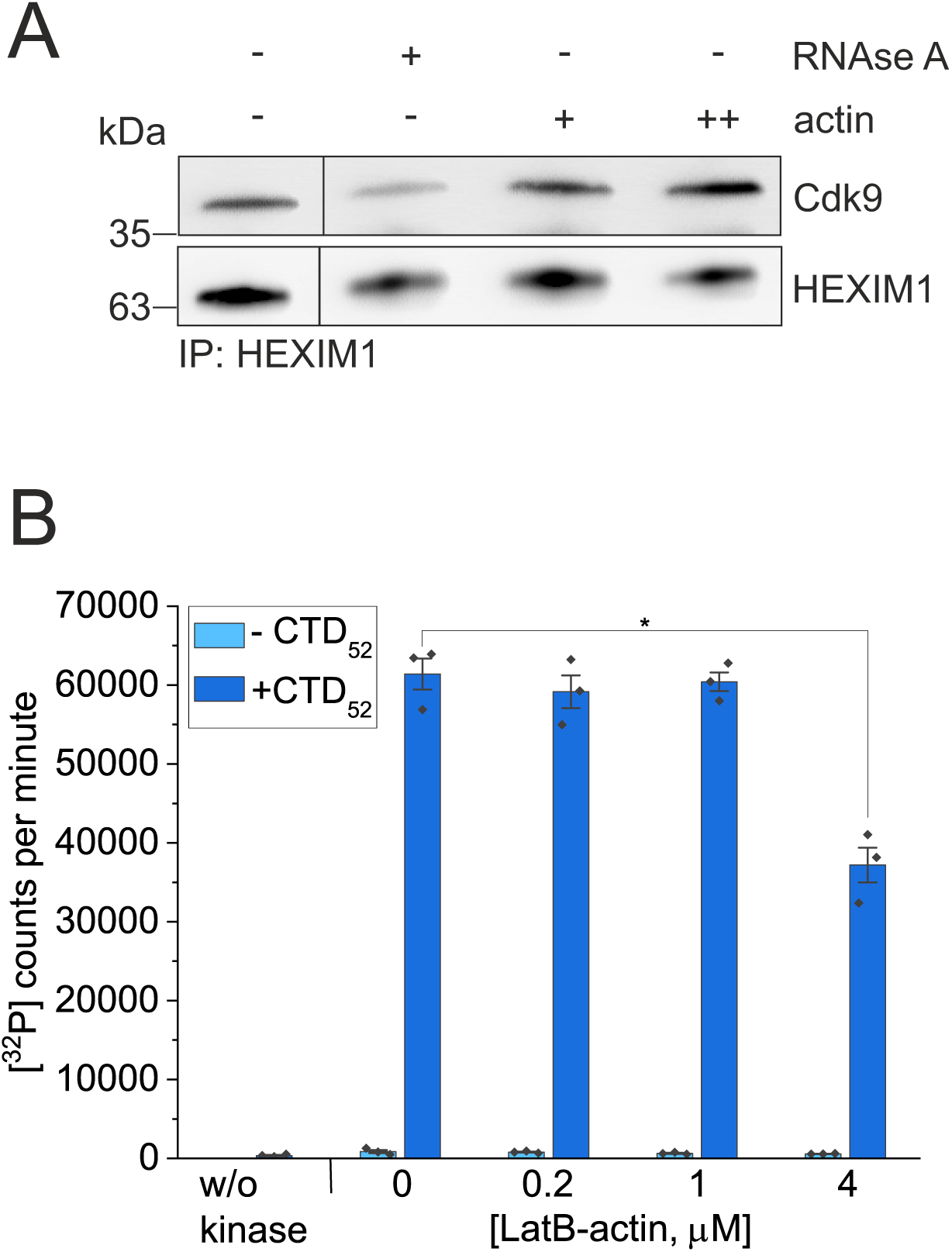
*In vitro*, actin does not influence P-TEFb activity. A. Western blot of P-TEFb release from HEXIM1 immunoprecipitation by RNAse A and increasing amounts of purified rabbit muscle actin (+ 0.5 μM and ++ 2 μM). P-TEFb probed with Cdk9 antibody and HEXIM1 with HEXIM1 antibody. In control sample (first lane), no release agent added. Boxed areas cut from the same exposure to remove an extra control sample. B. *In vitro* kinase assay assessing recombinant P-TEFb kinase activity for phosphorylation of CTD_52_ substrate, and without the substrate as a control, with increasing LatB-actin amounts added to the reaction. Mean of triplicate measurements is presented, error bars ± 0.5 SD. *Statistically significant differences (P<0.05) with a two-sample t-test for samples with CTD substrate: 0 vs 0.2 μM actin P=0.532, 0 vs. 1 μM actin P=0.730, 0 vs. 4 μM actin P=0.002.

Another intriguing possibility is that actin could regulate Cdk9 kinase activity. The function of P-TEFb in releasing paused Pol II is based on Cdk9 phosphorylating the negative transcription elongation factors NELF and DSIF and serines at position 2 in the Pol II C-terminal domain (CTD) (Fujinaga et al., 2023). We thus set out to do an *in vitro* kinase assay, in which the phosphorylation of a biologically relevant CTD substrate by P-TEFb in the absence and presence of LatB-actin was analyzed by monitoring radioactive ATP. Based on our results, actin does inhibit P-TEFb kinase activity, but only at high concentrations (Fig. 5B). Significant effect was observed only when actin was used at 20 times higher concentration than P-TEFb, in line with the relative affinity measured for the Cdk9-actin interaction (Fig 2D).

## Discussion

Actin was linked to RNA polymerase function already almost 40 years ago (Scheer et al., 1984). Since then, several studies have demonstrated defects in transcription upon actin manipulations, whether with antibodies against actin (Hofmann et al., 2004; Miyamoto et al., 2011; Philimonenko et al., 2004), by altering nucleo-cytoplasmic actin shuttling (Dopie et al., 2012; Knerr et al., 2023; Le et al., 2016; Sokolova et al., 2018; Wei et al., 2020), by actin mutants that are either predominantly monomeric or filamentous (Almuzzaini et al., 2016; Huang et al., 2022; Knerr et al., 2023; Qi et al., 2011; Serebryannyy et al., 2016; Wei et al., 2020; Ye et al., 2008) or by using different actin-binding drugs (McDonald et al., 2006; Miyamoto et al., 2011; Wu et al., 2006; Ye et al., 2008). Nevertheless, the molecular mechanisms by which actin affects RNA polymerase function have remained rather unclear. Here we use ChIP-seq and biochemical experiments with purified proteins to demonstrate that actin interacts with genes only upon their active transcription elongation and that actin binds directly to the Cdk9 kinase subunit of P-TEFb. Since P-TEFb plays an essential role in releasing paused Pol II to productive transcription elongation, we postulate that this interaction recruits actin to elongating polymerase to facilitate co-transcriptional processes.

Earlier studies on selected target genes had indicated variable chromatin-binding patterns for actin, with some studies finding actin only on promoter regions (Hu et al., 2004), some also on the gene coding regions (Obrdlik et al., 2008; Philimonenko et al., 2004), and yet others also on intragenic sites (Ye et al., 2008). Our ChIP-seq studies in *Drosophila* ovaries revealed that actin binds to the promoter region of essentially all transcribed genes, but on highly expressed genes, it is also found on the gene bodies (Sokolova et al., 2018). Thus, the chromatin-binding pattern of actin depends on the expression level of the gene. Nevertheless, our earlier study was not able to discern the exact step of transcription, where actin plays a role. Here we took advantage of the fact that actin is strongly associated with *hsp* genes (Fig 1C-E), which have been widely used to dissect transcriptional mechanisms due to their robust activation upon HS. We found that actin associates with *hsp* genes only upon their active transcription elongation, and is not significantly present on the genes in the absence of HS, although the RNA polymerase can be readily detected as the paused polymerase (Fig 1C). This indicated a functional role for actin specifically during the transcription elongation phase. However, we cannot exclude the possibility that actin could play a role also during other steps of transcription. For example, actin has also been linked to pre-initiation complex assembly (Hofmann et al., 2004). It is possible that actin in this complex is not accessible to the antibody utilized here, warranting development of further reagents to study chromatin-binding properties of different forms of actin.

Actin has been demonstrated to copurify with all three RNA polymerases (Egly et al., 1984; Fomproix and Percipalle, 2004; Hu et al., 2004; Smith et al., 1979). Nevertheless, it has not been detected as an integral polymerase subunit in any of the high-resolution structural studies that have recently been conducted e.g. using cryo-EM methods (Bernecky et al., 2016; Fianu et al., 2021; Liu et al., 2018). Accordingly, we did not detect actin in the highly pure Pol II prep (Fig S1A) and actin also did not bind to the purified Pol II in SEC experiments (Fig S1B). In this latter case, as well as in some other experiments in this study (Fig S1B, 2C, 5B), we had to use actin complexed with LatB to overcome the limitation of actin polymerizing in physiological salt conditions. Even though LatB is a small molecule polymerization inhibitor, it can prevent interactions taking place at the exact same position on actin. Latrunculins bind near the nucleotide binding cleft of actin (Furstner et al., 2007; Yarmola et al., 2000), which leaves the cleft between subdomains 1 and 3 where most actin binding proteins bind free for interactions (Dominguez, 2004). It is, however, possible that LatB prevents the interaction between actin and Pol II. Another intriguing idea is that actin could interact with Pol II as a filament. Indeed, Pol II clusters formed upon serum stimulation colocalize with nuclear actin filaments (Wei et al., 2020) and the links of nuclear myosins to Pol function have raised the idea of an acto-myosin motor operating during transcription (Percipalle and Vartiainen, 2019). Unfortunately, we were not able to test this biochemically, since the purified Pol II co-sedimented not only with actin filaments, but also with microtubules (data not shown).

Earlier studies with co-immunoprecipitation techniques (Qi et al., 2011) and our own nuclear actin interactome analysis (Viita et al., 2019) had shown an interaction between actin and P-TEFb, but not provided any biochemical details of this interaction. Here we significantly extend these studies by demonstrating that the interaction between actin and Cdk9, the kinase subunit of P-TEFb, is direct. Using purified proteins, the direct interaction was detected in a pulldown (Fig 2C), pyrene actin polymerization (Fig 3A-B) and NBD-actin monomer binding (Fig 2D-E) assays, and the specificity of the binding demonstrated e.g. by using other transcription regulating Cdk/cyclin complexes (Fig 3D-E). Unfortunately, our attempts to gain further biochemical and structural information of the actin-P-TEFb complex proved unsuccessful. It might be that the interaction is very dynamic, which would also be supported with the relatively low affinity measured from the NBD-actin assays (Fig 2D-E) and the fact in the nuclear actin interactome analysis, P-TEFb was detected as a high-confidence interactor only with BioID, which detects also transient interactions, and not with affinity purification, which requires stable complex formation (Viita et al., 2019). In addition, the interaction might be electrostatic, which leads to practical problems when performing the assays with actin and P-TEFb. When purified, P-TEFb is usually kept in a high salt buffer to preserve its integrity and kinase activity, whereas actin polymerizes in high salt conditions. It therefore turned out to be difficult to find experimental conditions, where the salt concentration is high enough to maintain P-TEFb functional, and actin would be kept monomeric. Interestingly, also Cdk12/Cyclin K complex inhibited nuclear actin polymerization in the pyrene actin polymerization assays (Fig 3D), but to lesser extent than Cdk9 or P-TEFb (Fig 3A-B), whereas Cdk7/Cyclin H/Mat1 did not influence pyrene actin polymerization (Fig 3E). Of note, Cdk12/Cyclin K and Cdk9/Cyclin T1 share structural similarity, having open conformations (Bosken et al., 2014), whereas Cdk7/Cyclin H form is more closed (Peissert et al., 2020). In the future, it will be interesting to study, if actin may play a role in regulating Cdk12/Cyclin K complex, and cellular processes dependent on it.

It is tempting to speculate that the interaction between actin and P-TEFb plays a role in recruiting this complex to elongating Pol II. Indeed, Qi et al. (2011) found that in nuclear extracts depleted of actin, Cdk9 was not recruited to immobilized transcriptional template, but the mechanism has remained unclear. In cells, most of P-TEFb is sequestered to the 7SK snRNP complex (C. Quaresma et al., 2016). Of note, one component of this complex, LARP7, was a hit in the BioID nuclear actin interactome assay (Viita et al., 2019). Thus, we speculated that actin could release P-TEFb from the 7SK snRNP, and thereby promote its recruitment to Pol II. However, we did not obtain evidence of this in the P-TEFb release assay with increasing actin concentrations (Fig 5A). Another intriguing possibility is that actin could influence P-TEFb by modulating the kinase activity of Cdk9. Qi et al. (2011) performed an *in vitro* kinase assay using immunoprecipitation of different actin mutants overexpressed in HeLa nuclear extracts, after which CTD template and ATP were added to the reaction and the results are followed by Western blot. In this experimental setup, G-actin, but not predominantly filamentous actin, was found to stimulate serine 2 phosphorylation of Pol II CTD. We tested effects of actin to P-TEFb kinase activity using purified proteins to avoid possible effects of other proteins in nuclear extracts. We found that contrary to earlier results, high concentration of actin leads to less kinase activity (Fig 5B). However, the effect was visible only at the highest actin concentration used. Therefore, it is not possible to conclude that in cells, actin would have a significant role in modulating the kinase activity of P-TEFb. It therefore seems that the stimulatory effect of actin on Cdk9 kinase activity detected earlier (Qi et al., 2011) is not direct, but mediated by another protein that remains to be identified.

Intriguingly, the kinase activity of Cdk9 is not required for its interaction with actin, since the interaction persisted in the presence of different Cdk9 inhibitors (Fig 4A). Moreover, actin accumulates to *Hsp70* gene upon heat shock even when flavopiridol is present (Fig 4B). It has been reported that although flavopiridol reduces Pol II CTD Ser2 phosphorylation, it does not affect Pol II or P-TEFb density on genes (Ni et al., 2004). This indicates that the kinase activity is not needed for recruitment of P-TEFb to the gene. Hence, we currently favor a model, where the direct binding to Cdk9 recruits actin to the elongating Pol II. There actin, through some additional binding partners, could stimulate Cdk9 activity as suggested earlier (Qi et al., 2011). This could explain why altering nuclear actin levels affects transcription elongation rate (Viita et al., 2019). In addition, actin could also recruit other nuclear complexes to facilitate co-transcriptional processes. For instance, earlier studies have demonstrated that actin interacts with several hnRNP proteins (Kukalev et al., 2005; Percipalle et al., 2002; Percipalle et al., 2001) and our nuclear actin interactome analysis identified numerous proteins implicated in RNA processing, including core components of the spliceosome (Viita et al., 2019). Detailed biochemical analysis of the underlying molecular interactions are needed to fully understand these tightly intertwined processes.

## Materials and methods

### Plasmids and cloning

pEF-Flag-G13R-actin and pEF-Flag-V159N-actin plasmids were a gift from Richard Treisman (The Francis Crick Institute, UK). 2HA-actin was cloned with restriction digestion. 2Flag-R62D-actin and 2Flag-G168D;Y169D-actin were cloned as described in Viita et al. 2019. pENTR221-Cdk9 plasmid was purchased from the Genome Biology Unit, University of Helsinki (clone number 100003812) and used to generate 2HA-Cdk9, 2Flag-Cdk9 and Cdk9-pDest10 baculoviral expression construct using pDest10 backbone from Thermo Fisher Scientific. His-Cdk9 (fl)/GST-Cyclin T1 (1-272), GST-Cdk7(2-346)/Cyclin H(1-323)/Mat1(1-309) and GST-Cdk12 (714-1063)/GST-CycK (1-267) in pACEBac1 expression constructs are described in Düster et al. 2022.

### Antibodies

The following antibodies were from Merck: Actin Ac-40 (A3853, 1:1000), Actin Ac-74 (A2228), HRP-conjugated polyHistidine (His-HRP, A7058, 1:5000), HRP-conjugated anti-goat IgG (A9452, 1:5000), HRP-conjugated anti-Flag M2 (A8592, 1:5000) and HRP-conjugated anti-HA (HA-7) (H6533, 1:5000). Rpb1 antibody (664906, 1:500) was from BioLegend and HEXIM1 from Everest Biotech (EB06964, 1:2000). Cdk9 antibody (2316S, 1:1000) was from Cell Signaling Technology. Anti-RNA polymerase II CTD repeat YSPTSPS (phospho S5) antibody (ab5408) was from Abcam. Normal mouse IgG antibody (sc-2025) was from Santa Cruz Biotechnology. The following antibodies were from Thermo Fisher Scientific: HRP-conjugated anti-rabbit IgG (G-21234, 1:5000), HRP-conjugated anti-mouse IgG (G-21040, 1:5000).

### Cell lines

HeLa cells obtained from Marikki Laiho (University of Helsinki) were cultured in DMEM High Glucose w Sodium Pyruvate w/o L-glutamine medium (EuroClone) supplemented with 10% fetal bovine serum (Thermo Fisher Scientific), penicillin-streptomycin (Thermo Fisher Scientific) and GlutaMAX (Thermo Fisher Scientific) in 5% CO_2_ incubator at +37°C. *Drosophila* Schneider cell line (S2R+) was obtained from Jussi Taipale (University of Helsinki) and cultured in Schneider’s *Drosophila* Medium (Thermo Fisher Scientific) supplemented with 10% heat inactivated fetal bovine serum (Thermo Fisher Scientific) and 1 x antibiotic-antimycotic solution (penicillin, streptomycin, Amphotericin B; Thermo Fisher Scientific) at +25°C incubator.

### Rabbit skeletal muscle actin purification and LatB-actin preparation

Actin was purified from rabbit muscle acetone powder (Pel-Freez Biologicals) as described in Pardee & Spudich 1982. LatB-actin was prepared by incubating rabbit skeletal muscle actin with a tenfold molar excess of LatB (Merck) overnight at +4°C. 20x initiation buffer (2 M NaCl, 60 mM MgCl_2_, 10 mM ATP) was added to polymerize uncomplexed actin and the possible filaments were removed by centrifugation at 200 000 g for 15 min at +4°C.

### RNA polymerase II purification

Pol II was purified from HeLa nuclear pellets as described in Kostek et al. 2006. Briefly, the nuclear pellets were solubilized to 50 mM Tris pH 7.9, 5 mM MgCl_2_, 0.5 mM EDTA, 25% glycerol and sonicated. An ammonium sulfate precipitation with a 42% cut was performed, the pellets were resuspended, dialyzed to 150 mM ammonium sulfate and placed on a DEAE52 anion exchange column (Whatman). After washing, Pol II was eluted to 400 mM ammonium sulfate, the protein containing fractions were pooled and dialyzed to a buffer with 200 mM ammonium sulfate. Next, Pol II was bound to protein G Sepharose Fast Flow beads (GE Healthcare) containing 8WG16 antibody against Rpb1, beads were washed with high salt and Pol II was eluted to a buffer with a tri-heptapeptide repeat of the CTD and concentrated.

### Recombinant protein expression and purification

His-Cdk9

His-Cdk9 was expressed in insect cells using the MultiBac expression system. Cdk9-pDest10 plasmid was transformed into DH10MultiBac cells, recombinant bacmids were prepared and the presence of correct construct was verified by PCR. The bacmid was transfected into Sf9 cells using Fugene HD transfection reagent (Promega). The obtained viral stock was amplified and used for protein expression. The expression of His-Cdk9 was done by infecting cells at the density of 2 x 10^6^ cells/ml with the baculoviral stock at a multiplicity of infection (MOI) of 1. After 96 hours, the cells were collected by centrifugation.

His-Cdk9 was purified by lysing the cell pellets in lysis buffer (25 mM Tris-HCl pH 8.0, 10 mM NaCl, 1 mM MgCl_2_, 2 mM β-mercaptoethanol, 0.2% Igepal) supplemented with cOmplete EDTA free protease inhibitor cocktail (Merck) and benzonase nuclease (Merck). Cell lysate was homogenized using EmulsiFlex-C3 (Avestin) with gauge pressure of 15 000 psi for 10 min. After homogenization, NaCl concentration of the lysate was increased to 200 mM and 20 mM imidazole was added. Cell extract was clarified by centrifugation at 66 000 g, 30 min at +4°C. Supernatant was applied to HisTrap HP 5 ml column (GE Healthcare) pre-equilibrated in 25 mM Tris-HCl pH 8.0, 300 mM NaCl, 2 mM β-mercaptoethanol, 20 mM imidazole buffer, column was washed with high salt (1 M NaCl) buffer and gradient elution using 500 mM imidazole was performed. Fractions containing protein of interest were analyzed by SDS-PAGE and Coomassie staining, pooled and injected to Superdex 200 HiLoad 16/600 gel filtration column (GE Healthcare) equilibrated with 25 mM Tris-HCl pH 7.5, 100 mM NaCl, 1 mM DTT, 0.5 mM EDTA, 10% glycerol buffer. Fractions were collected, protein containing fractions were analyzed and protein was concentrated and stored.

P-TEFb, Cdk7/CycH/Mat1, Cdk12/CycK expression

Recombinant proteins were expressed in Sf9 insect cells, except for P-TEFb cyclin subunit Cyclin T1 that was expressed in *E. coli* BL21(DE3). Cdk12/CycK were co-expressed with CAK1 from *Saccharomyces cerevisiae*. Expressions done as described in Düster et al. 2022.

P-TEFb purification

His-Cdk9 cell pellet was resuspended in lysis buffer (50 mM Hepes pH 7.6, 500 mM NaCl, 5 mM β-mercaptoethanol, 20 mM imidazole) supplemented with 1 mM PMSF and 1 μg/ml DNAseI. Lysate was sonicated, cleared by centrifugation and applied to 5 ml HisTrap column (GE Healthcare) pre-equilibrated in lysis buffer. Column was extensively washed with wash buffer (50 mM Hepes, 1 M NaCl, 5 mM β-mercaptoethanol) and protein was eluted with elution buffer (50 mM Hepes, 500 mM NaCl, 5 mM β-mercaptoethanol, 500 mM imidazole) by gradient elution. Protein containing fractions were pooled and processed to TEV protease cleavage (10 μg/ml) overnight to cleave His-tag.

GST-Cyclin T1 bacterial pellet was resuspended in lysis buffer (50 mM Hepes, 150 mM NaCl, 5 mM β-mercaptoethanol) supplemented with 1 mM PMSF and 1 μg/ml DNAseI. Lysate was sonicated, cleared by centrifugation and added to Pierce Glutathione Agarose beads (Thermo Scientific) pre-washed with lysis buffer. Protein was eluted with elution buffer (50 mM Hepes, 300 mM NaCl, 5 mM β-mercaptoethanol, 10 mM reduced glutathione) and eluted protein was processed to TEV protease cleavage overnight to cleave GST-tag.

Next day, proteins were centrifuged at 10 000 g, 10 min at +4°C to remove possible precipitates and combined in 1:1 molar ratio to reconstitute the P-TEFb complex. The protein mixture was then loaded to Superdex 200 HiLoad 16/600 column (GE Healthcare) equilibrated with SEC buffer (50 mM Hepes pH 7.6, 500 mM NaCl, 1 mM TCEP, 10% glycerol). Peak fractions containing equal amounts of Cdk9 and Cyclin T1 as monitored by SDS-PAGE were collected, concentrated and stored.

Cdk7/CycH/Mat1 purification

Purification described in Düster et al. 2022

Cdk12/CycK purification

Purification described in Bösken et al. 2014.

### Gel filtration

LatB-actin as a control (20 μM, 10 μl) or LatB-actin mixed with Pol II (2 μM, 10 μl) for one hour on rotation was run to Superose 6 Increase 3.2/300 column (GE Healthcare) connected to Pharmacia SMART chromatography system and equilibrated with G-actin buffer (5 mM Tris-HCl pH 8.0, 0.2 mM CaCl_2_, 0.2 mM ATP, 0.5 mM DTT) supplemented with 100 mM NaCl. Run was done at 0.06 ml/min and 120 μl fractions were collected until 1 column volume had passed. 4x SDS-PAGE loading buffer was added to fractions and they were processed to SDS-PAGE and Western blotting using actin Ac-40 and Pol II Rpb1 antibodies.

### Pulldown assay with purified proteins

His-GST lysate (10 μl) and purified His-Cdk9 (1 μM) in total volume of 300 μl were added to 50 μl of Ni-NTA agarose (Qiagen) equilibrated with binding buffer 1 (25 mM Tris-HCl pH 7.5, 150 mM NaCl, 1 mM DTT) and reactions were incubated for two hours on rotation at +4°C. Beads were washed once with binding buffer 1, three times with binding buffer 2 (25 mM Tris-HCl pH 7.5, 500 mM NaCl, 1 mM DTT) and once with binding buffer 1. LatB-actin (3 μM in total volume of 300 μl) was added to the beads and samples were incubated for four hours on rotation at +4°C. After incubation beads were washed three times with binding buffer 1 and bound proteins were eluted to 1x SDS-PAGE loading buffer and processed for SDS-PAGE and Western blotting using His-HRP, Cdk9 and actin Ac-40 antibodies.

### *In vitro* kinase assay

Radioactive kinase measurements were done as described previously (Duster et al., 2022). Recombinant, purified P-TEFb with full-length Cdk9 and short Cyclin T1 (1-272), 0.2 μM, was mixed with 0-4 μM LatB-actin concentrations in kinase buffer (50 mM Hepes, 34 mM KCl, 7 mM MgCl_2_, 2.5 mM DTE, 5 mM β-glycerol phosphate, pH 7.6). 0.2 mM final ATP concentration with 0.45 mCi [32P]-γ-ATP/ml were added and reactions were pre-incubated for 5 min. Reactions were then started by addition of CTD_52_ substrate (50 μM) and incubated for 15 min at 30°C in a shaking incubator at 300 rpm. Reactions were stopped by adding EDTA to a final concentration of 50 mM. Reaction mixtures were spotted onto filter sheets (Amersham Protran nitrocellulose membrane, GE Healthcare) and washed 3 x with PBS. Radioactive counts were assessed in a Beckman Liquid Scintillation Counter for 1 min. Measurements were conducted in triplicates and the mean ± 0.5 SD is presented.

### NBD-actin monomer binding assay

NBD-actin (ATP bound) was purchased from Cytoskeleton. To prepare ADP-actin, hexokinase beads were used (Pollard 1986). CNBr activated Sepharose 4B (GE Healthcare) were let to swell in 1 mM HCl pH 3.0 for 30 min and washed. Hexokinase (Merck) was diluted to coupling buffer (0.1 M NaHCO_3_ pH 8.3, 0.5 M NaCl), added to the beads and incubated overnight at +4°C. Next day, reactive groups were blocked with 0.1 M Tris-HCl pH 8.0 for one hour, beads were washed with 30 mM Tris-HCl pH 8.0, 1 mM CaCl_2_ and following materials were added: ATP-bound NBD-actin, 2.5 mM glucose, 0.4 mM ADP and 20 mM Tris-HCl pH 8.0. Beads were incubated for two hours at +4°C, after which ADP-NBD-actin was collected.

Measurement was done as described in Mattila et al. 2004. Reactions containing 10 x initiation mix (10 mM MgCl_2_, 1 M KCl), NBD-actin in final concentration of 0.2 μM and increasing concentrations of protein of interest in G-buffer (2 mM Tris-HCl pH 8, 0.1 mM CaCl_2_, 0.2 mM ATP or ADP, 0.1 mM DTT) were pipetted to a black 96-well plate and fluorescence was measured at 482/520 nm, 1 sec per well.

From the obtained data, the normalized enhancement of fluorescence was determined by the equation

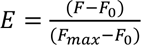

Where

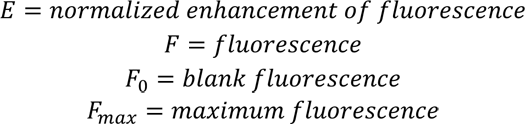

Data were analyzed and fitted using Origin 2022b and the following equation

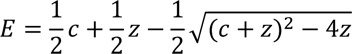

where z and c are determined in the two following equations

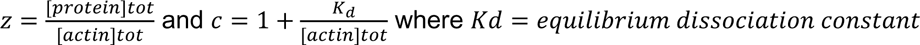

Relative fluorescence was presented as a function of [protein of interest]/[actin]. Curves were used to see if the binding saturates and efficiency of actin-binding was calculated.

### Pyrene actin polymerization assay

Pyrene actin (Cytoskeleton) and purified rabbit muscle actin were diluted to 20 μM in G-actin buffer (5 mM Tris-HCl pH 7.5, 0.2 mM CaCl_2_, 0.2 mM ATP, 0.2 mM DTT) and mixed to obtain 5% pyrene labeled actin. Mixture was centrifuged at 312 500 g for 30 min at +4°C to ensure completely monomeric actin at the starting point of measurement. Reactions were prepared by diluting protein of interest to G-buffer and adding 10 x KME buffer (5 mM Tris-HCl pH 7.5, 1 M KCl, 50 mM MgCl_2_, 10 mM EGTA) that creates conditions for actin polymerization. For pyrene actin assay with Cdk9 inhibitors, the inhibitors were also added at this point. Finally, actin was added to the reaction, reaction was transferred immediately to a cuvette and fluorescence measurement was started to catch the starting point of actin polymerization. Measurement was done with sampling period of 10 sec, illumination 5 nm, detection 10 nm, wavelengths 365 nm/407 nm, total time 4000 sec. Data shown by plotting the measured absorbance as a function of time.

### Co-immunoprecipitation (co-IP)

HeLa cells were plated on 10 cm cell culture dishes at a density of 1 x 10^6^ cells per plate. Next day, cells were co-transfected with HA and Flag-tagged constructs (3 + 3 μg) using JetPrime transfection reagent (Polyplus) according to the manufacturer’s protocol. After two days of transfection, cells were resuspended to cold IP-buffer (50 mM Tris-HCl pH 7.5, 150 mM NaCl, 0.5% Igepal and cOmplete EDTA free protease inhibitor cocktail (Merck)) and lysate was cleared by centrifugation. Pre-washed anti-Flag M2 affinity gel (Merck) or EZview Red Anti-HA Affinity Gel (Merck), 25 μl of slurry per sample, was added to lysates and samples were incubated on rotation at +4°C for four hours. After incubation, samples were washed three times with IP buffer without Igepal and bound proteins were eluted with 1x SDS-PAGE loading buffer and processed for SDS-PAGE and Western blotting using HA-HRP and Flag-HRP antibodies.

### P-TEFb release assay

P-TEFb release assay was performed as described in Bugai et al. 2019. Briefly, whole cell extract from one confluent 15 cm plate of HeLa cells was prepared by lysing the cells to 0.5 ml of buffer C (20 mM Tris-HCl, 150 mM NaCl, 10 mM KCl, 1.5 mM MgCl_2_, 0.5 mM EDTA, 10% glycerol, 0.5% Igepal, pH 7.9) supplemented with cOmplete EDTA free protease inhibitor cocktail and incubating for one hour on rotation at +4°C. After incubation, lysate was cleared by centrifugation. To immobilize 7SK snRNP, 1:2 diluted whole cell extract in buffer C was added to Dynabeads Protein G (Thermo Fisher Scientific) pre-incubated with HEXIM1 antibody, using 2 μg antibody and 20 μl protein G solution per sample, and incubated on rotation at +4°C for four hours. After incubation, beads were washed once with buffer C containing 350 mM NaCl, once with buffer C and once with G-buffer (5 mM Tris-HCl pH 8.0, 0.2 mM CaCl_2_, 0.2 mM ATP, 0.5 mM DTT). For release reactions, either 5 μl PureLink RNAse A (Thermo Fisher Scientific) or actin in increasing concentrations in G-buffer was added to the beads containing immobilized 7SK snRNP, reactions were incubated on rotation at +4°C for two hours, beads were washed once with G-buffer and once with buffer C and 1x SDS-PAGE loading buffer was added to the beads to elute bound proteins. Samples were processed to SDS-PAGE and Western blotting using HEXIM1 antibody to see equal amounts of HEXIM1 and Cdk9 antibody to observe possible release from 7SK snRNP.

### ChIP

S2R+ cells were splitted in T75 culture flasks and for ChIP-qPCR treated with 300 nM flavopiridol (MedChemExpress) or equal volume of DMSO for 10 min. Heat shock was performed by placing sealed flasks into +37°C water bath for 20 mins. Samples were crosslinked with 1% formaldehyde for 10 min and crosslinking was quenched with ice-cold 125 mM glycine on ice for 5 min. Chromatin fragments were obtained from cells lysed in RIPA buffer (10 mM Tris-HCl pH 8.0, 140 mM NaCl, 1 mM EDTA, 1% Triton X-100, 0.1% SDS, 0.1% sodium deoxycholate) by sonication with Diagenode Bioruptor, 30 sec on and 30 sec off with the highest power setting, for 15 cycles in case of ChIP-seq and for 8 cycles in ChIP-qPCR. Cells from one T75 flask were used for each immunoprecipitation. Immunoprecipitation was carried out with 5 μg of antibody (IgG, actin Ac-74 or Pol II S5P) overnight at +4°C and immune-complexes were collected with 50 μl of protein A sepharose (GE Healthcare) at +4°C for two hours on rotation. Chromatin was eluted from washed beads in 1% SDS in TE buffer (10 mM Tris-HCl pH 8.0, 1 mM EDTA). DNA purification after reverse crosslinking was done with phenol/chloroform/isoamyl alcohol (Thermo Fisher Scientific) and DNA was precipitated with 0.1 volume of 3 M sodium acetate pH 5.2 and two volumes of ethanol using glycogen as a carrier.

### ChIP-seq

ChIP libraries were prepared using NEBNext ChIP-seq DNA Sample Prep Master Mix Set for Illumina (NEB E6240) and NEBNext Multiplex Oligos for Illumina (Index Primers Set 1, NEB E7335) according to manufacturer’s instructions. Sequencing was done with Illumina NextSeq500 at Biomedicum Functional Genomics Unit (FuGU). ChIP-seq was performed in duplicates. ChIP-seq data sets were aligned using Bowtie2 (using Chipster software; Kallio et al. 2011) to version dm6 of *Drosophila* genome with the default settings. Genome coverage files were generated using DeepTools with RPKM normalization. To analyze, visualize and present ChIP-seq data, we used Integrative Genomics Viewer (IGV, Robinson et al. 2011) and DeepTools.

### Data availability

S2R+ ChIP-seq datasets generated in this manuscript have been deposited to Gene Expression Omnibus under accession GSE239771. ChIP-seq files corresponding to *Drosophila* w^1118^ ovaries were downloaded from GEO under accession number GSE116362.

### ChIP-qPCR

ChIP-qPCR was performed using SensiFAST SYBR No-ROX One-Step Kit (Meridian Bioscience) according to manufacturer’s instructions using 1:10 diluted inputs and ChIP samples without diluting.

Primers for *Drosophila* Hsp70 middle region (Hsp70mid), 5’ to 3’ orientation:

Forward: CGAGATTGACGCACTGTTTG

Reverse: CACGATGTCGTGGATCTGAC

Percent of input calculated noting the dilution factor of input, using the equation

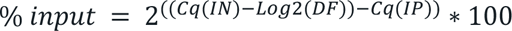

### Statistical analyses

Statistical testing was performed using Origin 2022b. Normality was tested and when data conformed to a normal distribution, two-tailed two-sample t-test was used. All statistical tests were done with the significance level of 0.05.

## Acknowledgements

We acknowledge Paula Maanselkä for excellent technical assistance throughout this project. Sequencing was performed at Biomedicum Functional Genomics Unit (FuGU) supported by Helsinki Institute of Life Science (HiLIFE) and Biocenter Finland at the University of Helsinki and Novogene. Fly food was provided by HiFLY, supported by HiLIFE and Biocenter Finland. This work was supported by Sigrid Juselius, Cancer as well as Jane and Aatos Erkko foundations as well as Academy of Finland grants 338281 and 330254 to MKV and Instrumentarium Science Foundation and EMBO Scientific Exchange Grant 9864 to SK.

---

**Figure S1.**
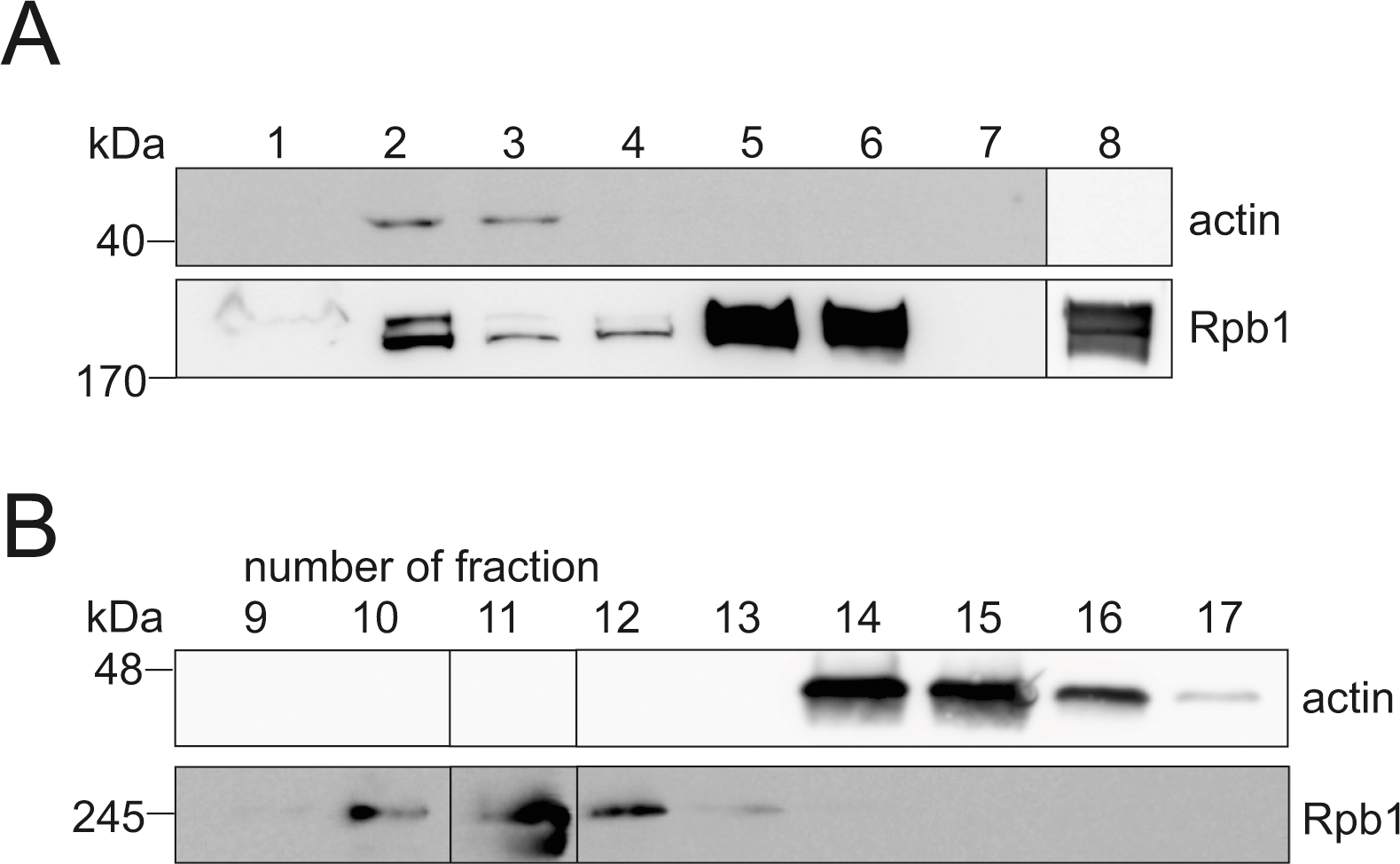
Actin does not interact with the core RNA polymerase II. A. Western blot of actin (Ac-40) and Pol II subunit Rpb1 from samples taken during the Pol II purification process (1-7) and the purified polymerase (8). Boxed areas are from separate Western blots. 1. sample during ammonium sulfate precipitation, 2. sample after ammonium sulfate precipitation, 3. flow-through from anion exchange column, 4. wash of anion exchange column, 5. sample after anion exchange, 6. flow-through from affinity purification, 7. wash of affinity beads, 8. final purified Pol II. B. Western blot of gel filtration with LatB-actin and Pol II fractions using Ac-40 antibody for actin and Rpb1 antibody for Pol II. Boxed areas are cut from the same exposure to present samples loaded to two gels in a continuous manner.

## References

Almuzzaini B, Sarshad AA, Rahmanto AS, Hansson ML, Von Euler A, Sangfelt O, Visa N, Farrants AK and Percipalle P (2016) In beta-actin knockouts, epigenetic reprogramming and rDNA transcription inactivation lead to growth and proliferation defects. FASEB J 30:2860–2873.

Ambrosio L and Schedl P (1984) Gene expression during Drosophila melanogaster oogenesis: analysis by in situ hybridization to tissue sections. Dev Biol 105:80–92.

Baumli S, Lolli G, Lowe ED, Troiani S, Rusconi L, Bullock AN, Debreczeni JE, Knapp S and Johnson LN (2008) The structure of P-TEFb (CDK9/cyclin T1), its complex with flavopiridol and regulation by phosphorylation. EMBO J 27:1907–1918.

Bernecky C, Herzog F, Baumeister W, Plitzko JM and Cramer P (2016) Structure of transcribing mammalian RNA polymerase II. Nature 529:551–554.

Bieniasz PD, Grdina TA, Bogerd HP and Cullen BR (1999) Recruitment of cyclin T1/P-TEFb to an HIV type 1 long terminal repeat promoter proximal RNA target is both necessary and sufficient for full activation of transcription. Proc Natl Acad Sci U S A 96:7791–7796.

Blanchoin L and Pollard TD (1998) Interaction of actin monomers with Acanthamoeba actophorin (ADF/cofilin) and profilin. J Biol Chem 273:25106–25111.

Bosken CA, Farnung L, Hintermair C, Merzel Schachter M, Vogel-Bachmayr K, Blazek D, Anand K, Fisher RP, Eick D and Geyer M (2014) The structure and substrate specificity of human Cdk12/Cyclin K. Nat Commun 5:3505.

Bugai A, Quaresma AJC, Friedel CC, Lenasi T, Duster R, Sibley CR, Fujinaga K, Kukanja P, Hennig T, Blasius M, Geyer M, Ule J, Dolken L and Barboric M (2019) P-TEFb Activation by RBM7 Shapes a Pro-survival Transcriptional Response to Genotoxic Stress. Mol Cell 74:254–267 e210.

C. Quaresma A, Bugai A and Barboric M (2016) Cracking the control of RNA polymerase II elongation by 7SK snRNP and P-TEFb. Nucleic Acids Res 44:7527–7539.

Cao T, Sun L, Jiang Y, Huang S, Wang J and Chen Z (2016) Crystal structure of a nuclear actin ternary complex. Proc Natl Acad Sci U S A 113:8985–8990.

Carlier MF, Jean C, Rieger KJ, Lenfant M and Pantaloni D (1993) Modulation of the interaction between G-actin and thymosin beta 4 by the ATP/ADP ratio: possible implication in the regulation of actin dynamics. Proc Natl Acad Sci U S A 90:5034–5038.

Carlier MF, Laurent V, Santolini J, Melki R, Didry D, Xia GX, Hong Y, Chua NH and Pantaloni D (1997) Actin depolymerizing factor (ADF/cofilin) enhances the rate of filament turnover: implication in actin-based motility. J Cell Biol 136:1307–1322.

Chao SH, Fujinaga K, Marion JE, Taube R, Sausville EA, Senderowicz AM, Peterlin BM and Price DH (2000) Flavopiridol inhibits P-TEFb and blocks HIV-1 replication. J Biol Chem 275:28345–28348.

Core L and Adelman K (2019) Promoter-proximal pausing of RNA polymerase II: a nexus of gene regulation. Genes Dev 33:960–982.

Detmers P, Weber A, Elzinga M and Stephens RE (1981) 7-Chloro-4-nitrobenzeno-2-oxa-1,3-diazole actin as a probe for actin polymerization. J Biol Chem 256:99–105.

Dominguez R (2004) Actin-binding proteins--a unifying hypothesis. Trends Biochem Sci 29:572–578.

Doolittle LK, Rosen MK and Padrick SB (2013) Measurement and analysis of in vitro actin polymerization. Methods Mol Biol 1046:273–293.

Dopie J, Skarp KP, Kaisa Rajakyla E, Tanhuanpaa K and Vartiainen MK (2012) Active maintenance of nuclear actin by importin 9 supports transcription. Proc Natl Acad Sci U S A 109:E544–552.

Duster R, Ji Y, Pan KT, Urlaub H and Geyer M (2022) Functional characterization of the human Cdk10/Cyclin Q complex. Open Biol 12:210381.

Egly JM, Miyamoto NG, Moncollin V and Chambon P (1984) Is actin a transcription initiation factor for RNA polymerase B? EMBO J 3:2363–2371.

Fianu I, Dienemann C, Aibara S, Schilbach S and Cramer P (2021) Cryo-EM structure of mammalian RNA polymerase II in complex with human RPAP2. Commun Biol 4:606.

Fiore A, Spencer VA, Mori H, Carvalho HF, Bissell MJ and Bruni-Cardoso A (2017) Laminin-111 and the Level of Nuclear Actin Regulate Epithelial Quiescence via Exportin-6. Cell Rep 19:2102–2115.

Fomproix N and Percipalle P (2004) An actin-myosin complex on actively transcribing genes. Exp Cell Res 294:140–148.

Fujinaga K, Huang F and Peterlin BM (2023) P-TEFb: The master regulator of transcription elongation. Mol Cell 83:393–403.

Furstner A, De Souza D, Turet L, Fenster MD, Parra-Rapado L, Wirtz C, Mynott R and Lehmann CW (2007) Total syntheses of the actin-binding macrolides latrunculin A, B, C, M, S and 16-epi-latrunculin B. Chemistry 13:115–134.

Hofmann WA, Stojiljkovic L, Fuchsova B, Vargas GM, Mavrommatis E, Philimonenko V, Kysela K, Goodrich JA, Lessard JL, Hope TJ, Hozak P and de Lanerolle P (2004) Actin is part of pre-initiation complexes and is necessary for transcription by RNA polymerase II. Nat Cell Biol 6:1094–1101.

Hu P, Wu S and Hernandez N (2004) A role for beta-actin in RNA polymerase III transcription. Genes Dev 18:3010–3015.

Huang Q, Wu D, Zhao J, Yan Z, Chen L, Guo S, Wang D, Yuan C, Wang Y, Liu X and Xing J (2022) TFAM loss induces nuclear actin assembly upon mDia2 malonylation to promote liver cancer metastasis. EMBO J 41:e110324.

Kallio MA, Tuimala JT, Hupponen T, Klemela P, Gentile M, Scheinin I, Koski M, Kaki J and Korpelainen EI (2011) Chipster: user-friendly analysis software for microarray and other high-throughput data. BMC Genomics 12:507.

Klages-Mundt NL, Kumar A, Zhang Y, Kapoor P and Shen X (2018) The Nature of Actin-Family Proteins in Chromatin-Modifying Complexes. Front Genet 9:398.

Knerr J, Werner R, Schwan C, Wang H, Gebhardt P, Grotsch H, Caliebe A, Spielmann M, Holterhus PM, Grosse R and Hornig NC (2023) Formin-mediated nuclear actin at androgen receptors promotes transcription. Nature 617:616–622.

Kostek SA, Grob P, De Carlo S, Lipscomb JS, Garczarek F and Nogales E (2006) Molecular architecture and conformational flexibility of human RNA polymerase II. Structure 14:1691–1700.

Kukalev A, Nord Y, Palmberg C, Bergman T and Percipalle P (2005) Actin and hnRNP U cooperate for productive transcription by RNA polymerase II. Nat Struct Mol Biol 12:238–244.

Le HQ, Ghatak S, Yeung CY, Tellkamp F, Gunschmann C, Dieterich C, Yeroslaviz A, Habermann B, Pombo A, Niessen CM and Wickstrom SA (2016) Mechanical regulation of transcription controls Polycomb-mediated gene silencing during lineage commitment. Nat Cell Biol 18:864–875.

Liu X, Farnung L, Wigge C and Cramer P (2018) Cryo-EM structure of a mammalian RNA polymerase II elongation complex inhibited by alpha-amanitin. J Biol Chem 293:7189–7194.

Lu H, Xue Y, Yu GK, Arias C, Lin J, Fong S, Faure M, Weisburd B, Ji X, Mercier A, Sutton J, Luo K, Gao Z and Zhou Q (2015) Compensatory induction of MYC expression by sustained CDK9 inhibition via a BRD4-dependent mechanism. eLife 4:e06535.

Mattila PK, Quintero-Monzon O, Kugler J, Moseley JB, Almo SC, Lappalainen P and Goode BL (2004) A high-affinity interaction with ADP-actin monomers underlies the mechanism and in vivo function of Srv2/cyclase-associated protein. MolBiolCell 15:5158–5171.

Mattila PK, Salminen M, Yamashiro T and Lappalainen P (2003) Mouse MIM, a tissue-specific regulator of cytoskeletal dynamics, interacts with ATP-actin monomers through its C-terminal WH2 domain. JBiolChem 278:8452–8459.

McDonald D, Carrero G, Andrin C, de Vries G and Hendzel MJ (2006) Nucleoplasmic beta-actin exists in a dynamic equilibrium between low-mobility polymeric species and rapidly diffusing populations. J Cell Biol 172:541–552.

Miyamoto K, Pasque V, Jullien J and Gurdon JB (2011) Nuclear actin polymerization is required for transcriptional reprogramming of Oct4 by oocytes. Genes Dev 25:946–958.

Ni Z, Schwartz BE, Werner J, Suarez JR and Lis JT (2004) Coordination of transcription, RNA processing, and surveillance by P-TEFb kinase on heat shock genes. Mol Cell 13:55–65.

Obrdlik A, Kukalev A, Louvet E, Farrants AK, Caputo L and Percipalle P (2008) The histone acetyltransferase PCAF associates with actin and hnRNP U for RNA polymerase II transcription. Mol Cell Biol 28:6342–6357.

Ojala PJ, Paavilainen VO, Vartiainen MK, Tuma R, Weeds AG and Lappalainen P (2002) The two ADF-H domains of twinfilin play functionally distinct roles in interactions with actin monomers. MolBiolCell 13:3811–3821.

Olson CM, Jiang B, Erb MA, Liang Y, Doctor ZM, Zhang Z, Zhang T, Kwiatkowski N, Boukhali M, Green JL, Haas W, Nomanbhoy T, Fischer ES, Young RA, Bradner JE, Winter GE and Gray NS (2018) Pharmacological perturbation of CDK9 using selective CDK9 inhibition or degradation. Nat Chem Biol 14:163–170.

Pardee JD and Spudich JA (1982) Purification of muscle actin. Methods Cell Biol 24:271–289.

Peissert S, Schlosser A, Kendel R, Kuper J and Kisker C (2020) Structural basis for CDK7 activation by MAT1 and Cyclin H. Proc Natl Acad Sci U S A 117:26739–26748.

Percipalle P, Jonsson A, Nashchekin D, Karlsson C, Bergman T, Guialis A and Daneholt B (2002) Nuclear actin is associated with a specific subset of hnRNP A/B-type proteins. Nucleic Acids Res 30:1725–1734.

Percipalle P and Vartiainen M (2019) Cytoskeletal proteins in the cell nucleus: a special nuclear actin perspective. Mol Biol Cell 30:1781–1785.

Percipalle P, Zhao J, Pope B, Weeds A, Lindberg U and Daneholt B (2001) Actin bound to the heterogeneous nuclear ribonucleoprotein hrp36 is associated with Balbiani ring mRNA from the gene to polysomes. J Cell Biol 153:229–235.

Philimonenko VV, Zhao J, Iben S, Dingova H, Kysela K, Kahle M, Zentgraf H, Hofmann WA, de Lanerolle P, Hozak P and Grummt I (2004) Nuclear actin and myosin I are required for RNA polymerase I transcription. Nat Cell Biol 6:1165–1172.

Posern G, Sotiropoulos A and Treisman R (2002) Mutant actins demonstrate a role for unpolymerized actin in control of transcription by serum response factor. Mol Biol Cell 13:4167–4178.

Qi T, Tang W, Wang L, Zhai L, Guo L and Zeng X (2011) G-actin participates in RNA polymerase II-dependent transcription elongation by recruiting positive transcription elongation factor b (P-TEFb). J Biol Chem 286:15171–15181.

Ramirez F, Ryan DP, Gruning B, Bhardwaj V, Kilpert F, Richter AS, Heyne S, Dundar F and Manke T (2016) deepTools2: a next generation web server for deep-sequencing data analysis. Nucleic Acids Res 44:W160–165.

Robinson JT, Thorvaldsdottir H, Winckler W, Guttman M, Lander ES, Getz G and Mesirov JP (2011) Integrative genomics viewer. Nat Biotechnol 29:24–26.

Scheer U, Hinssen H, Franke WW and Jockusch BM (1984) Microinjection of Actin-Binding Proteins and Actin Antibodies Demonstrates Involvement of Nuclear Actin in Transcription of Lampbrush Chromosomes. Cell 39:111–122.

Serebryannyy LA, Parilla M, Annibale P, Cruz CM, Laster K, Gratton E, Kudryashov D, Kosak ST, Gottardi CJ and de Lanerolle P (2016) Persistent nuclear actin filaments inhibit transcription by RNA polymerase II. J Cell Sci 129:3412–3425.

Sidorenko E and Vartiainen MK (2019) Nucleoskeletal regulation of transcription: Actin on MRTF. Exp Biol Med (Maywood) 244:1372–1381.

Smith SS, Kelly KH and Jockusch BM (1979) Actin co-purifies with RNA polymerase II. BiochemBiophysResCommun 86:161–166.

Sokolova M, Moore HM, Prajapati B, Dopie J, Merilainen L, Honkanen M, Matos RC, Poukkula M, Hietakangas V and Vartiainen MK (2018) Nuclear Actin Is Required for Transcription during Drosophila Oogenesis. iScience 9:63–70.

Spencer VA, Costes S, Inman JL, Xu R, Chen J, Hendzel MJ and Bissell MJ (2011) Depletion of nuclear actin is a key mediator of quiescence in epithelial cells. J Cell Sci 124:123–132.

Stuven T, Hartmann E and Gorlich D (2003) Exportin 6: a novel nuclear export receptor that is specific for profilin.actin complexes. EMBO J 22:5928–5940.

Ulferts S, Prajapati B, Grosse R and Vartiainen MK (2021) Emerging Properties and Functions of Actin and Actin Filaments Inside the Nucleus. Cold Spring Harb Perspect Biol 13.

Vartiainen MK, Guettler S, Larijani B and Treisman R (2007) Nuclear actin regulates dynamic subcellular localization and activity of the SRF cofactor MAL. Science 316:1749–1752.

Vartiainen MK, Mustonen T, Mattila PK, Ojala PJ, Thesleff I, Partanen J and Lappalainen P (2002) The three mouse actin-depolymerizing factor/cofilins evolved to fulfill cell-type-specific requirements for actin dynamics. Mol Biol Cell 13:183–194.

Viita T, Kyheroinen S, Prajapati B, Virtanen J, Frilander MJ, Varjosalo M and Vartiainen MK (2019) Nuclear actin interactome analysis links actin to KAT14 histone acetyl transferase and mRNA splicing. J Cell Sci.

Vinson VK, De La Cruz EM, Higgs HN and Pollard TD (1998) Interactions of Acanthamoeba profilin with actin and nucleotides bound to actin. Biochemistry 37:10871–10880.

Wei M, Fan X, Ding M, Li R, Shao S, Hou Y, Meng S, Tang F, Li C and Sun Y (2020) Nuclear actin regulates inducible transcription by enhancing RNA polymerase II clustering. Sci Adv 6:eaay6515.

Wu X, Yoo Y, Okuhama NN, Tucker PW, Liu G and Guan JL (2006) Regulation of RNA-polymerase-II-dependent transcription by N-WASP and its nuclear-binding partners. Nat Cell Biol 8:756–763.

Yarmola EG, Somasundaram T, Boring TA, Spector I and Bubb MR (2000) Actin-latrunculin A structure and function. Differential modulation of actin-binding protein function by latrunculin A. J Biol Chem 275:28120–28127.

Ye J, Zhao J, Hoffmann-Rohrer U and Grummt I (2008) Nuclear myosin I acts in concert with polymeric actin to drive RNA polymerase I transcription. Genes Dev 22:322–330.

Zheng X, Diraviyam K and Sept D (2007) Nucleotide effects on the structure and dynamics of actin. Biophys J 93:1277–1283.

